# Targeting the ITGA3B1 Complex with A Novel Antibody-Drug Conjugate for Metastatic Bladder Cancer

**DOI:** 10.1101/2025.04.06.647354

**Authors:** Hyeryeon Jung, Junghyeon Lee, Eunhee G. Kim, Jinsung Ahn, Michael C. Haffner, Peter S. Nelson, Kwang Pyo Kim, John K. Lee, Eugene C. Yi, Kristine M. Kim

**Affiliations:** Department of Molecular Medicine and Biopharmaceutical Sciences, Graduate School of Convergence Science and Technology, Seoul National University, Seoul 03080, Republic of Korea; Division of Human Biology, Fred Hutchinson Cancer Center, Seattle, WA 98119; Department of Bio-Health Convergence, Kangwon National University, Chuncheon 24341, Republic of Korea; Department of Systems Immunology, Division of Biomedical Convergence, College of Biomedical Science, Kangwon National University, Chuncheon 24341, Republic of Korea; Department of Laboratory Medicine and Pathology, University of Washington, Seattle, WA 98195; Department of Applied Chemistry, Department of Applied Chemistry, Institute of Natural Science, Kyung Hee University, Yongin, Republic of Korea; Department of Biomedical Science and Technology, Kyung Hee Medical Science Research Institute, Kyung Hee University, Seoul, Republic of Korea; Division of Hematology/Oncology, Department of Medicine, David Geffen School of Medicine at UCLA, Los Angeles, CA 90095; Parker Institute for Cancer Immunotherapy at UCLA, David Geffen School of Medicine at UCLA, Los Angeles, CA 90095

## Abstract

Advances in antibody (Ab) phage display technologies have enabled the development of targeted therapies with enhanced precision and clinical efficacy. Here, we report a phenotypic screening strategy combining target-unbiased live-cell biopanning with in situ chemical crosslinking and mass spectrometry to identify internalizing antibodies and their cognate antigens—key features for effective antibody-drug conjugate (ADC) development. Using this approach, we identified the 2E7 antibody, which exhibits rapid internalization and high specificity for the integrin α3β1 (ITGA3B1) heterodimer, a complex overexpressed in multiple solid tumors, including bladder cancer. We generated an ITGA3B1-targeted ADC by conjugating monomethyl auristatin E (MMAE) to the 2E7 antibody, enabling selective delivery of cytotoxic payloads to ITGA3B1-positive cells. In preclinical bladder cancer models, this ADC demonstrated potent and dose-dependent antitumor efficacy, with significant tumor regression and improved survival. Our findings establish a framework for target discovery using live-cell phenotypic screening and position ITGA3B1 as a promising therapeutic target for ADC-based treatment of advanced bladder cancer.

**Teaser:** Live-cell biopanning uncovers ITGA3B1 as a target for ADC development in metastatic bladder cancer.

## Introduction

Antibodies (Abs) that target cell-surface proteins have reshaped the landscape of cancer therapy by enabling precise diagnostic and therapeutic interventions (1, 2). Among the strategies driving this transformation, live-cell Ab selection has emerged as a powerful approach that overcomes limitations of conventional methods reliant on purified recombinant proteins, which often lack native conformation and post-translational modifications. By screening directly in the context of intact, living cells, this technique identifies Abs that recognize antigens in their physiologic state, preserving native structural complexity and functional relevance (3-6).

Unlike traditional Ab discovery platforms that focus primarily on affinity for pre-selected targets, live-cell panning enables phenotypic selection based on cellular behavior, viability, and morphology (7). This unbiased strategy allows for the simultaneous identification of Abs and their corresponding antigens without prior knowledge of target identity (8). Moreover, screening in a physiologically relevant cellular environment improves the predictive value for therapeutic efficacy, particularly in challenging settings such as primary and stem-like cancer cells (9). Critically, live-cell platforms also enrich for antibodies with internalization capacity—an essential feature for antibody-drug conjugates (ADCs), which rely on intracellular delivery of cytotoxic payloads (10-13).

Live-cell Ab discovery has demonstrated therapeutic potential across multiple indications. For example, cell-based platforms have identified novel targets in ovarian cancer (14) and yielded Abs with improved efficacy in chronic lymphocytic leukemia compared to rituximab (15). However, the complexity of the cell-surface proteome—where key targets may be low-abundance or embedded in dynamic protein complexes—poses a major barrier to antigen deconvolution (16). To overcome this, we developed an integrative in situ crosslinking mass spectrometry strategy to stabilize and isolate native antibody-antigen complexes, enabling accurate identification of cognate targets. Such strategies have been effective in uncovering novel ligand-receptor interactions, including the identification of B7H6 as a ligand for the Nkp30 receptor in immune regulation (17).

Using this platform, we identified the 2E7 Ab, which rapidly internalizes and selectively binds the integrin α3β1 (ITGA3B1) complex—a heterodimeric receptor implicated in tumor invasion and metastasis. ITGA3B1 is highly expressed in several solid tumors, including bladder cancer, where it correlates with aggressive disease phenotypes (18-20). We conjugated 2E7 to monomethyl auristatin E (MMAE), a potent antimitotic agent, to generate a targeted ADC. In preclinical models of bladder cancer, this ADC induced robust, dose-dependent tumor regression and prolonged survival, with minimal toxicity.

Together, these findings underscore the value of phenotypic live-cell biopanning for identifying internalizing Abs and validating complex, tumor-specific targets. They further establish ITGA3B1 as a promising candidate for therapeutic intervention and demonstrate the feasibility of translating phenotypically selected Abs into effective ADCs for the treatment of advanced solid tumors.

## Results

### Live cell biopanning for identification of internalizing antibodies

We employed a differential live cell biopanning platform to identify Abs with efficient internalization capabilities specific to target cancer cells, as depicted in Figure 1A. We have previously established that phage Ab’s internalization can be effectively monitored during the biopanning process (11). Our findings confirm that the rates and levels of internalization of scFv-phage Abs correlate positively with their scFv and scFv-Fc forms, thus facilitating early determination of Ab internalization capabilities. We enriched a pool of phage-Abs specific to cancer stem cells derived from MDA-MB453 cells (CSCs-MDA-MB453) (21. 22) incorporating depletion (negative selection) step against CHO-DG44 cells and PBMCs, followed by three rounds of biopanning. After a 2-hour incubation at 37°C to induce internalization, the enriched phage-Abs were expanded in E. coli. A panel of phage Ab clones was then isolated and screened for binding to both CSCs-MDA-MB453 and CHO-DG44 cells. Seven clones (2E1, 2E11, 2H7, 3C3, 2E7, 2H6, and 3C7) were selected, converted into their scFv-Fc forms as previously described (5), and further evaluated for target cell specificity using flow cytometry against MDA-MB453 cells (Figure 1B).

**Figure 1.**
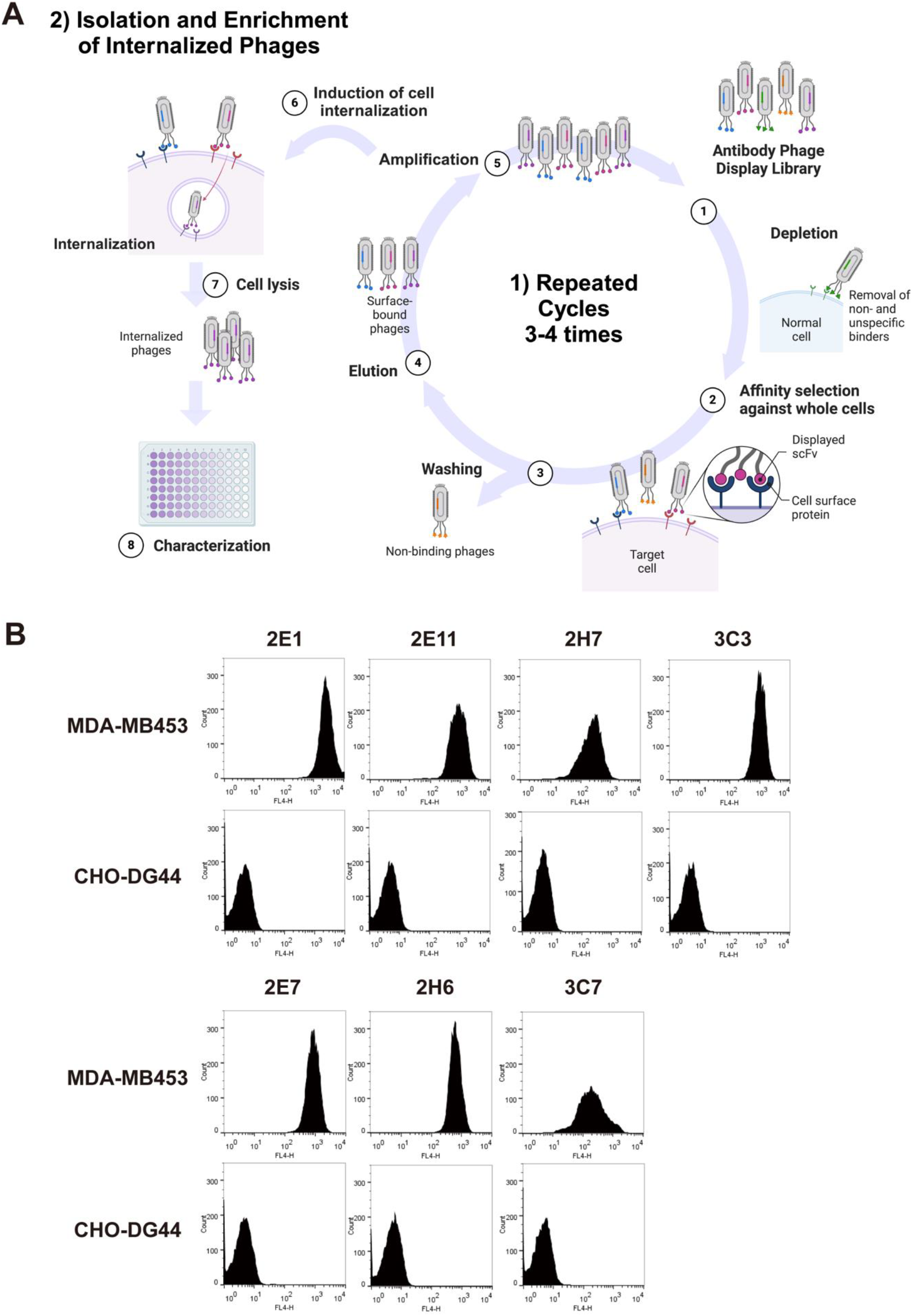
Schematic diagram of the live cell phage-display panning cycle for discovering cellular internalizing antibodies. **(A)**This diagram outlines the sequential steps involved in the phage-display panning process, utilized to isolate Abs that are capable of internalization into target cells. The process includes: 1) Incubation of a depleted phage-antibody (phage-Ab) library with target cells to enable specific binding to cell surface antigens. 2-3) Removal of unbound phages through washing. 4-6) Elution of both cell surface-bound and internalized phages. 7-8) The eluted phages are then recovered and amplified in E. coli TG1 for further analysis and characterization. **(B)** The target cell specificity of seven Ab clones isolated from the phage-display panning. Specificity is demonstrated through binding affinity to cell-surface proteins on MDA-MB-453 cancer cells compared to the control cell line, CHO-DG44.

Using flow cytometry and confocal microscopy, we extensively analyzed the internalization dynamics of the 2E7 scFv-Fc Ab in MDA-MB453 cells. Initial time-course flow cytometry results, as shown in Figure 2A, demonstrated efficient internalization rates with clones 2E1 and 2E7 reaching up to 89.2 ± 3.3% and 86.5 ± 2.5% internalization, respectively, within 90 minutes. Furthermore, the internalization kinetics of both the phage-Ab and their scFv-Fc derivatives were found to be similar, with the scFv-Fc forms of 2E1 and 2E7 reaching internalization rates of 94.4 ± 1.3% and 85.6 ± 2.1%, respectively, as shown in Figure 2B. In contrast, the other clones displayed internalization rates ranging from 20 to 60% at the same time point. Complementing these findings, confocal microscopy provided visual confirmation of the Ab’s behavior at the cellular level. Initially visible on the cell membrane as indicated by green fluorescence, the 2E7 scFv-Fc Ab underwent rapid uptake within 2 hours at 37°C, transitioning to overlapping orange fluorescence, which indicates localization within the lysosomal compartment (Figure 2C). This consistent pattern across single confocal sections confirms that the Ab is internalized through endocytosis and directed to intracellular compartments. These visualization studies support the quantitative data from flow cytometry, emphasizing the 2E7 scFv-Fc Ab effective navigation through cellular pathways to reach intracellular targets. The precise localization to lysosomes suggests the potential of the 2E7 scFv-Fc Ab for payload delivery directly to these organelles. This capability is particularly valuable in cancer treatment, where efficient internalization into target cells is crucial for therapeutic efficacy.

**Figure 2.**
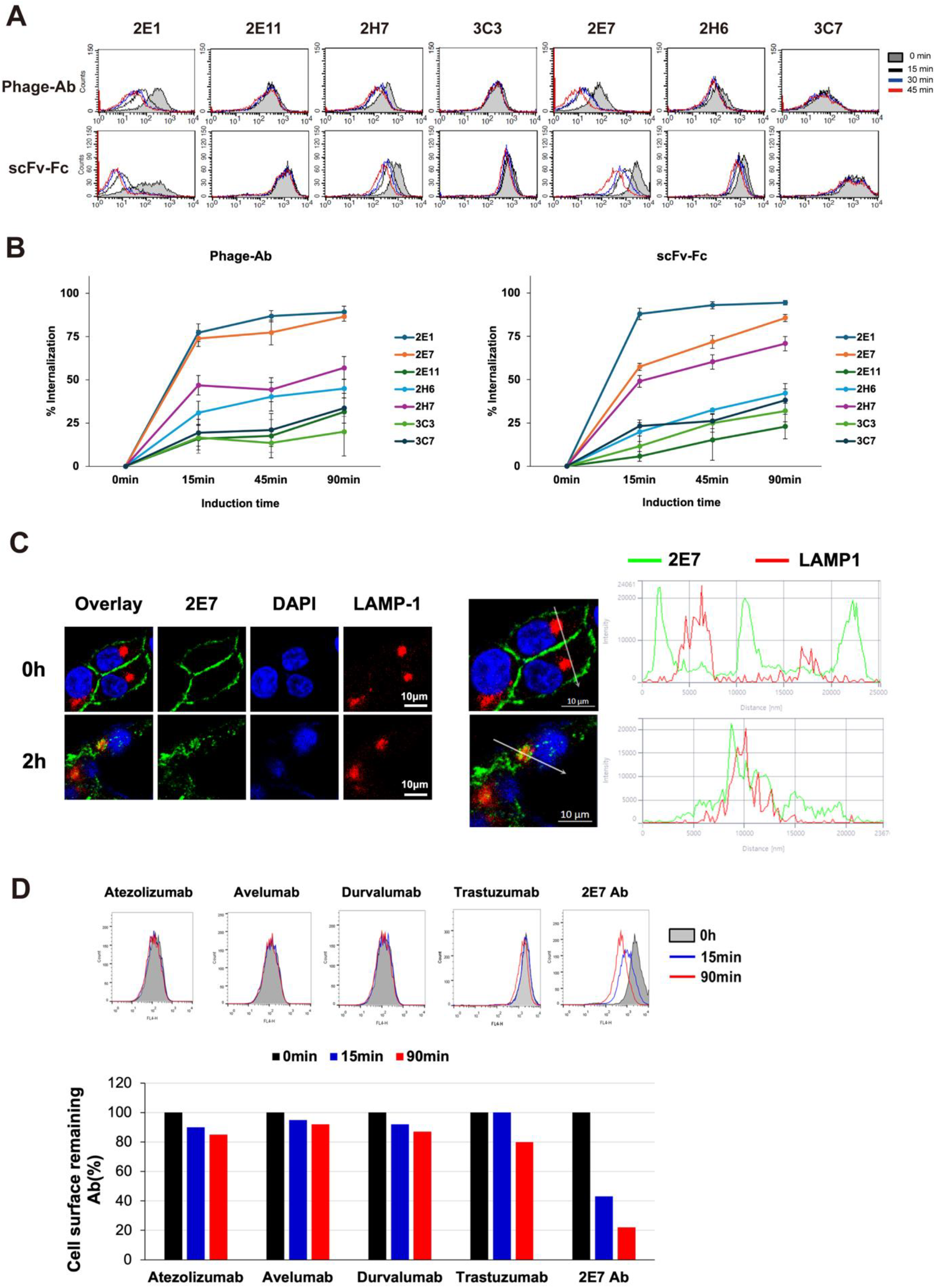
Evaluation of 2E7 Ab Internalization via FACS and Confocal Microscopy. **(A, B)** Flow Cytometry-Based Internalization Assay: Internalization of the 2E7 scFv-Fc Ab was quantified at designated time points. Abs were used at a concentration of 25 nM. Data represent geometric mean ± SD from three independent experiments, indicating the extent of internalization. **(C)** Confocal Microscopy of Ab Internalization: Cells were incubated with 2E7 scFv-Fc Abs and subsequently imaged at 0 hours (100% control) and 2 hours post-treatment at 37°C. Images show the co-localization of the scFv-Fc Abs (green) with the lysosomal marker LAMP-1 (red), and nuclei stained in blue. An enlarged view from the yellow highlighted area shows co-localization within the lysosomal compartment. Fluorescence intensity plots along the indicated arrows demonstrate overlapping signals of scFv-Fc Ab and LAMP-1, confirming their co-localization. **(D)** Internalization rates of 2E7 scFv-Fc Ab, Trastuzumab, and PD-L1 Abs: Flow cytometry and confocal microscopy were used to assess internalization of 2E7 scFv-Fc Ab, Trastuzumab, and PD-L1 antibodies (Atezolizumab, Avelumab, Durvalumab) in MDA-MB453 cells after 90 minutes at 37°C. Confocal microscopy confirmed 2E7 scFv-Fc Ab localization to lysosomes. Data are shown as mean internalization percentage ± SD (n=3).

We compared the internalization dynamics of the 2E7 scFv-Fc Ab with Her-2 Ab (Trastuzumab) and three FDA-approved PD-L1 Abs (Atezolizumab, Avelumab, and Durvalumab) commonly used in ADC generation. Under identical conditions, the 2E7 scFv-Fc Ab demonstrated significantly higher internalization rates, while Trastuzumab exhibited only 20% internalization, and the PD-L1 antibodies showed negligible internalization within 90 minutes (Figure 2D). This enhanced internalization capability of 2E7 scFv-Fc Ab highlights its potential for more efficient cytotoxic agent delivery in cancer treatment, suggesting superior therapeutic efficacy compared to other antibodies. These results, corroborated by both quantitative flow cytometry and qualitative confocal microscopy, emphasize the 2E7 scFv-Fc Ab’s ability to efficiently navigate cellular pathways and localize to lysosomal compartments, a key feature for effective ADC applications. This comprehensive analysis positions 2E7 scFv-Fc Ab as a promising candidate for targeted cancer therapy, particularly in cases requiring rapid and effective intracellular delivery.

### Target antigen identification for the 2E7 scFv-Fc Ab using in situ crosslinking, immunoprecipitation, and LC-MS/MS analysis

To identify the antigen specifically recognized by the 2E7 Ab clone, we utilized in situ chemical crosslinking combined with mass spectrometry analysis (18). This method provided a detailed analysis of the antigen specifically bound by the 2E7 Ab clone. As illustrated in Figure 3A, the scFv-Fc form of the 2E7 Ab clone was incubated with both cancer stem cells (CSCs) and non-cancer stem cells (NCSCs) in the presence of the amine-reactive chemical crosslinker BS3 s(bis(sulfosuccinimidyl)suberate), with human immunoglobulin G (HuIgG) serving as a negative control. Following cell lysis, antibody-antigen complexes were immunoprecipitated using Protein A magnetic beads and subsequently resolved by SDS-PAGE (Figure 3B) and then analyzed by LS-MS/MS and protein database search to identify the specific antigen. Western blot analysis confirmed the presence of crosslinked complexes in the high molecular weight region only in samples from the 2E7 Ab clone with CSCs, not in those with NCSCs or the HuIgG control, indicating a specific recognition of an antigen present uniquely in CSCs (Figure 3C).

**Figure 3.**
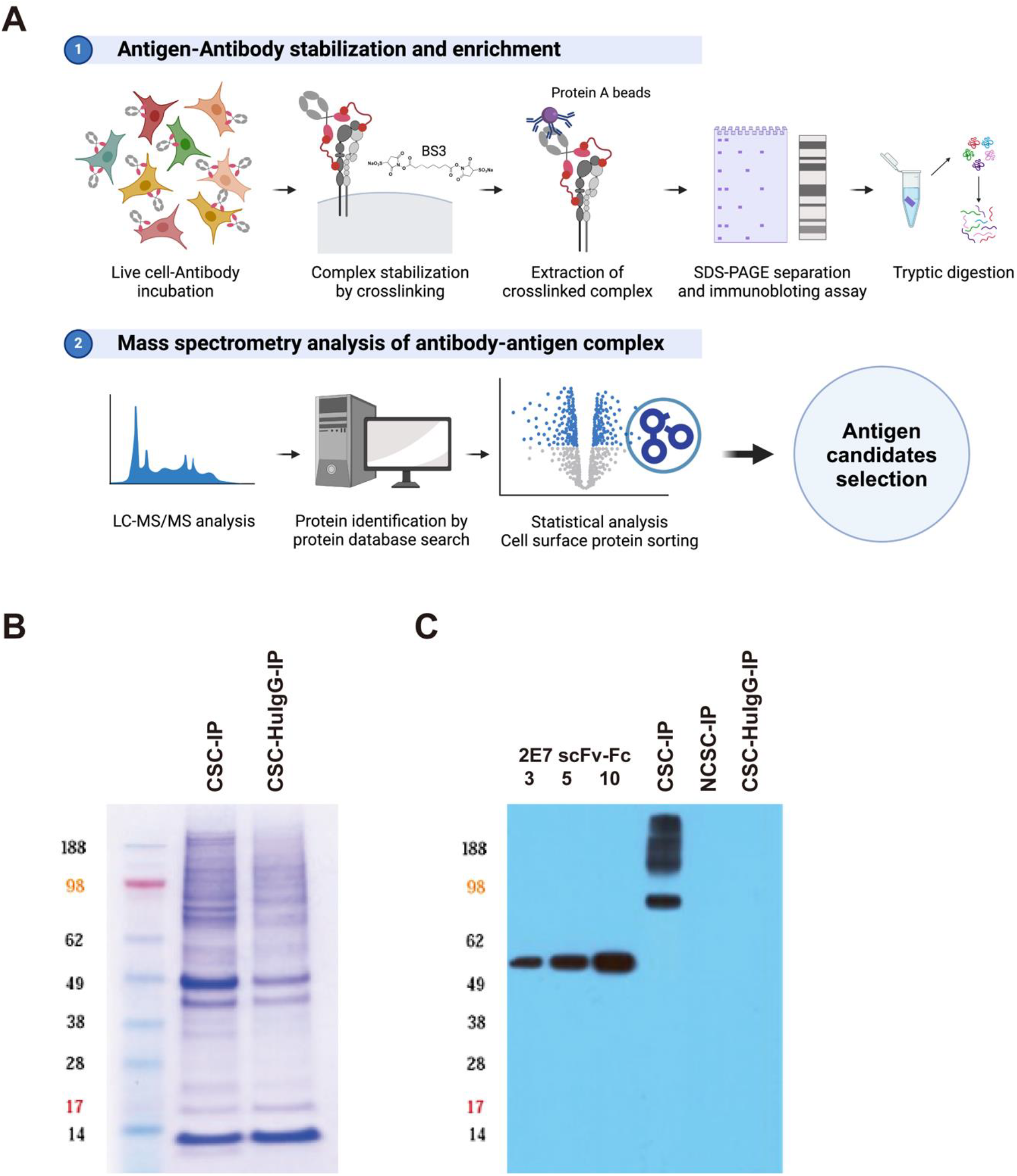
Identification of the target antigen for the 2E7 scFv-Fc Ab using LC-MS/MS analysis. **(A)** Schematic overview of the methodology employed for the target antigen identification, detailing each step from sample preparation to LC-MS/MS analysis and protein database search. **(B)** Representative SDS-PAGE gel illustrating the separation of proteins within the crosslinked complex after immunoprecipitation of the 2E7 scFv-Fc Ab with CSCs and NCSCs. **(C)** Western blot analysis confirmed the specific detection of the target antigen within the crosslinked complex from the 2E7 scFv-Fc Ab, showing no crosslinked products in the NCSCs and huIgG control samples.

For MS analysis, in-gel tryptic digestion was performed on the corresponding high molecular weight gel region (>80 kDa) for both IP samples from the 2E7 scFv-Fc and the HuIgG. LC-MS/MS analyses and subsequent protein database searches using Peaks Studio (ver. 8.5) identified a total of 837 proteins after excluding keratin contaminants, each supported by two or more unique peptides (STN > 2.0, FDR < 0.01%) (23). After excluding proteins identified in the HuIgG crosslinked complexes, we focused on 40 potential antigens for the 2E7 scFv-Fc Ab, selecting only plasma membrane based on Gene Ontology Cellular Component (GOCC) analysis. We detected subsets of integrin protein with high confidence (−10lgP p-value >20) enriched in the 2E7 scFv-Fc crosslinked complexes. These subsets comprised five integrin alpha subunit proteins (ITGA3, ITGA6, ITGA2, and ITGAV) and two are integrin beta subunits (ITGB1 and ITGB4). These were identified with the highest spectral counts in the 2E7 scFv-Fc crosslinked complexes, significantly exceeding those observed in the negative control sample (PLGEM, STN >2.0. p-value <0.01), as shown in Table 1 (24). Known for their ability to form heterodimers in situ, as detailed in Table 1, these integrins prompted further investigation into specific alpha-beta pairings (25-27). Our subsequent verification experiments focused on the interaction between integrin alpha-3 (ITGA3) and beta-1 (ITGB1), which are known to form the ITGA3B1 complex. This pairing is particularly significant, as ITGA3 is uniquely known to complex with ITGB1, highlighting a specific interaction critical for cellular functions (25).

**Table 1.** Identified subsets of Integrin proteins enriched in the 2E7 scFv-Fc crosslinked complexes.

PLGEM, Power Law Global Error Model; STN: Signal To Noise
| Accession number | Description | Gene Name | -10lgP | PLGEM |  | Integrin group (26) | Heterodimer (27) |
| --- | --- | --- | --- | --- | --- | --- | --- |
|  |  |  |  | STN | p-value |  |  |
| P05556 | Integrin beta-1 | ITGB1 | 326.48 | 15.02 | 0.00 |  | A1, A2, A10, A11, A9, A4, A5, AV, A8, A3, A6, A7 |
| P16144 | Integrin beta-4 | ITGB4 | 421.64 | 13.82 | 0.00 |  | A6 |
| P26006 | Integrin alpha-3 | ITGA3 | 355.81 | 13.64 | 0.00 | Laminin receptors | B1 |
| P23229 | Integrin alpha-6 | ITGA6 | 327.24 | 6.92 | 0.00 | Laminin receptors | B1, B4 |
| P17301 | Integrin alpha-2 | ITGA2 | 235.82 | 2.84 | 0.00 | Collagen receptors | B1 |
| P06756 | Integrin alpha-V | ITGAV | 152.29 | 2.34 | 0.01 | Fibronectin receptors | B1, B5, B6 |

### Validation of 2E7 scFv-Fc Ab binding specificity to ITGA3B1 heterodimer complexes

Building upon our initial immunoprecipitation-mass spectrometry (IP-MS) findings, we conducted ELISA analyses to validate the specificity of the 2E7 scFv-Fc Ab toward the heterodimeric complexes of integrin alpha and beta subunits, specifically ITGA3B1 and ITGA6B4. Using integrin complexes ITGA3B1 and ITGA6B4, along with their respective subunits ITGA3, ITGB1, ITGA6, and ITGB4, we assessed the binding affinities of the 2E7 scFv-Fc Ab. huIgG and BSA were employed as controls for non-specific binding, enabling us to precisely delineate the specific interactions of 2E7 scFv-Fc Ab. The Ab demonstrated specific and enhanced binding to the ITGA3B1 complex over its individual subunits or the ITGA6B4 complex (Figure 4A). Notably, negligible binding was observed to ITGA6 and ITGB4 subunits, confirming the high specificity of 2E7 for ITGA3B1. Further concentration-dependent binding assay revealed an EC50 of 1.2 nM for ITGA3 alone, improving to 0.9 nM when bound to the ITGA3B1 heterodimer. This indicates a preference for the functional heterodimer complex (Figure 4B).

**Figure 4.**
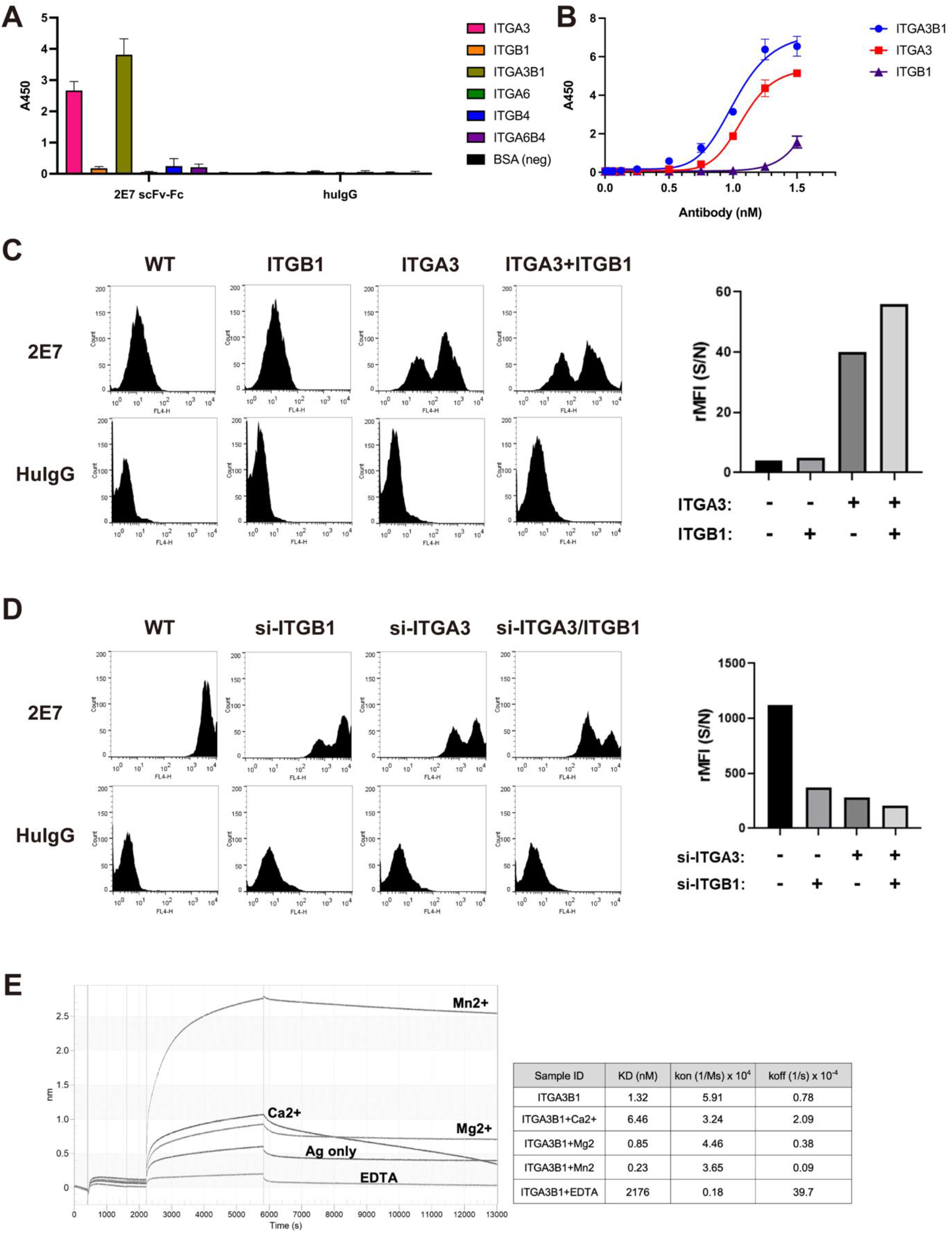
In vitro and in vivo binding specificity of the 2E7 scFv-Fc Ab. **(A)** ELISA analysis of 2E7 scFv-Fc binding to ITGA3B1 vs. isolated subunits and ITGA6B4. **(B)** Concentration-dependent binding of 2E7 scFv-Fc to ITGA3 and ITGA3B1. **(C)** Binding assay results in HEK293 cells transfected with ITGA3, ITGB1, or both. **(D)** Validation of binding specificity in MDA-MB231 cells with targeted knockdown of ITGA3 and ITGB1. **(E)** SPR analysis showing metal cofactor dependency of 2E7 scFv-Fc binding to ITGA3B1.

To further validate these findings in a physiologically relevant cell-based model, we employed HEK293 cells, which have low endogenous expression of ITGA3, for a binding assay. Cells were transiently transfected with plasmids expressing ITGA3, ITGB1, or both, to model the native conformation of the ITGA3B1 complex. The 2E7 Ab demonstrated significant binding enhancement exclusively in cells co-expressing ITGA3 and ITGB1, consistent with our ELISA results (Figure 4C). Negligible binding was observed in non-transfected or ITGB1-only transfected cells, further confirming the Ab’s specificity. For additional validation, we used the MDA-MB231 cell line, which naturally expresses high levels of both ITGA3 and ITGB1. Sequential knockdown of ITGB1, ITGA3, or both subunits via siRNA resulted in progressively decreased binding in ITGB1-knockdown, ITGA3-knockdown, and double-knockdown cells (Figure 4D). This gradient of binding affinity supports the conclusion that the 2E7 scFv-Fc Ab specifically targets the ITGA3/ITGB1 heterodimeric complex.

Furthermore, metal cofactor-dependency assays have deepened our understanding of the 2E7 scFv-Fc Ab’s binding characteristics. Surface plasmon resonance (SPR) biosensor analysis revealed that the interaction between the Ab and the ITGA3B1 complex is significantly enhanced by the presence of divalent metal cations such as Mg^2+^, and Mn^2+^, compared to the antigen alone. Notably, the presence of EDTA completely abolished binding, highlighting the critical role of these cations in facilitating the interaction (Figure 4E). Among the cations tested, Mn2+ produced the greatest enhancement, achieving a dissociation constant (kD) of 0.23 nM, compared to 6.5 nM with Ca^2+^ and 0.85 nM with Mg^2+^. These findings highlight the essential role of metal ions in stabilizing the active conformation of the ITGA3B1 complex (29), thereby significantly improving the antibody’s affinity and potential therapeutic effectiveness.

### Generation and characterization of MMAE conjugated 2E7 scFv-Fc ADC

The 2E7 scFv-Fc Ab targeting ITGA3B1 (anti-ITGA3B1 Ab) has demonstrated superior cellular internalization characteristics with target specificity, as described, supporting its potentiality as effective “driver” for cytotoxic drug compounds in an ADC (30). To explore the therapeutic potential of anti-ITGA3B1 Ab as a novel ADC, we conjugated with the highly potent microtubule inhibitor monomethyl auristatin E (MMAE) to the reduced interchain cysteines of the ITGA3B1 Ab using a maleimide-containing linker (Figure 5A) (31, 32). After conjugation, the size-exclusion chromatography (SEC) analysis showed that approximately 98% of the MMAE-conjugated 2E7 scFv-Fc was monomeric (Figure S1), suggesting that MMAE conjugation did not significantly affect the physiochemical properties of the antibody. Using a reversed-phase (RP)-HPLC method, MMAE conjugated anti-ITGA3B1 ADC resolved primarily into 4 and 6 drug to antibody ratio (DAR) species with an average DAR value of 4.09 (Figure 5B). Each peak was assigned its DAR by MALDI-TOF analysis (Figure S2).

**Figure 5.**
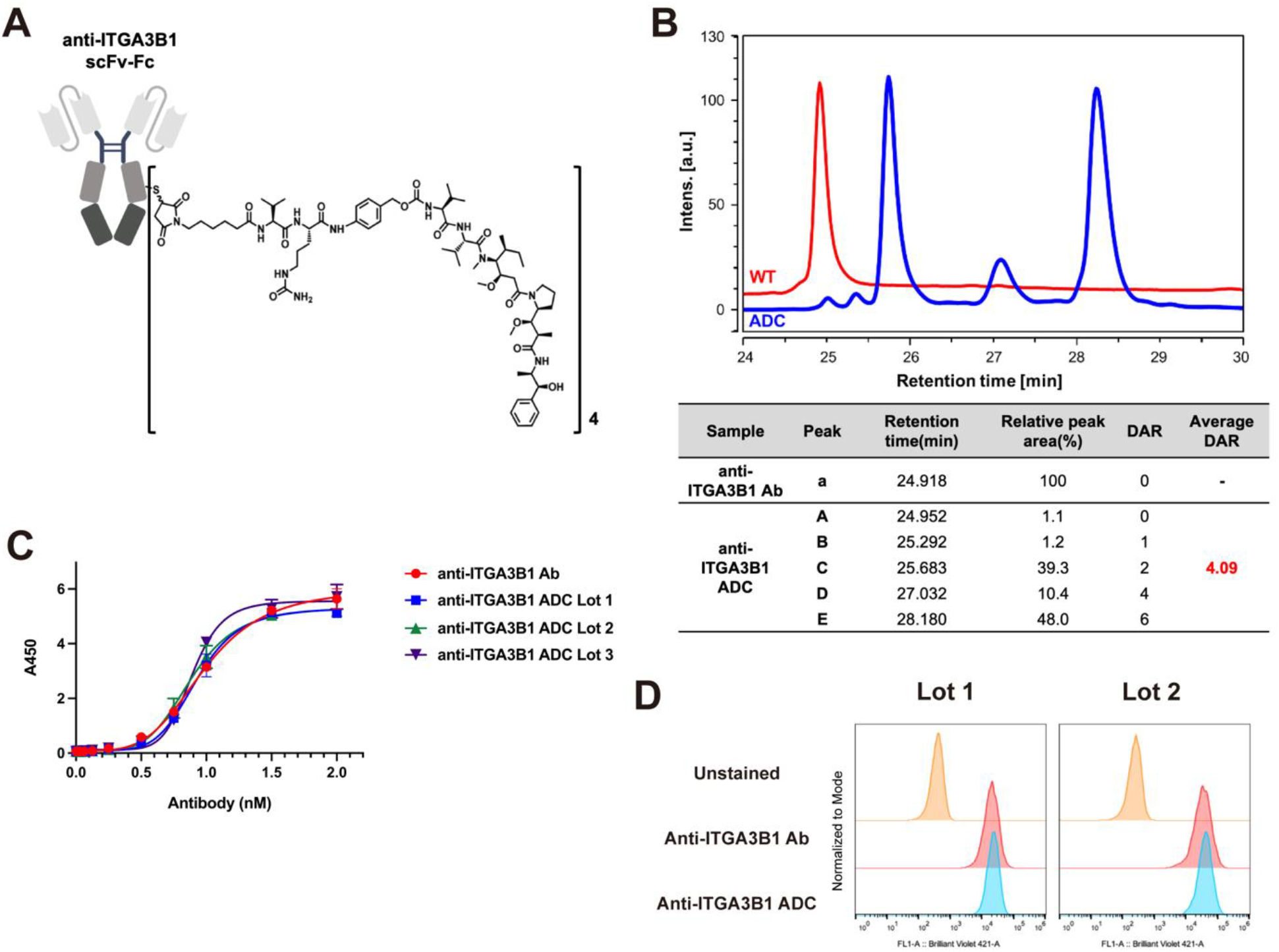
Characterization of MMAE-ITGA3B1 ADC. **(A)** Schematic diagram of vcMMAE-ITA3B1 ADC. **(B)** Overlapped RPLC chromatographic elution profiles of MMAE-ITGA3B1 ADCs (0-6 DAR). **(C)** ELISA antigen binding assays comparing three different lots of conjugated and an unconjugated ITGA3B1. **(D)** Flow cytometry analysis of UM-UC-3 cells stained with unconjugated and conjugated ITGA3B1. The red histograms represent staining with the corresponding unconjugated Ab, the blue histograms represent staining with the conjugated Ab, and yellow histograms indicate autofluorescence from unstained cells.

Additionally, we evaluated the binding retainability of both the conjugated and unconjugated anti-ITGA3B1 Abs via in vitro and in vivo binding assays. The unconjugated and three different lots of conjugated anti-ITGA3B1 Abs retained similar binding affinities towards ITGA3B1 in an Ab concentration-dependent manner (Figure 5C). Furthermore, flow cytometry analysis was conducted to compare the binding affinity of the conjugated and unconjugated Abs towards the ITGA3B1 high expression cell line. Both forms exhibited similar binding affinity to the cell line (Figure 5D), indicating that MMAE conjugation did not adversely affect the Ab’s binding properties.

### ITGA3 and ITGB1 Expression Profiling in Bladder Cancer

To examine ITGA3 expression profiles in bladder cancer, we utilized data from The Cancer Genome Atlas (TCGA) (33). This analysis revealed significant upregulation of ITGA3 not only bladder cancer but also in other human carcinomas, such as glioblastoma and pancreatic cancer, as detailed in Supplementary Figure 3. Specifically, Figure 6A shows that ITGA3 levels are substantially higher in bladder cancer patients (n = 408) compared to normal controls (n=18), highlighting a pronounced expression difference within a large dataset. Further analysis of ITGA3 across phenotypic subtypes of bladder cancer indicated a significant elevation in the basal squamous subtype, known for its aggressive behavior and poor prognosis (34), as depicted in Figure 6B. This observation supports the targeted application of our anti-ITGA3B1 Ab in therapeutic strategies for this challenging cancer subtype.

**Figure 6.**
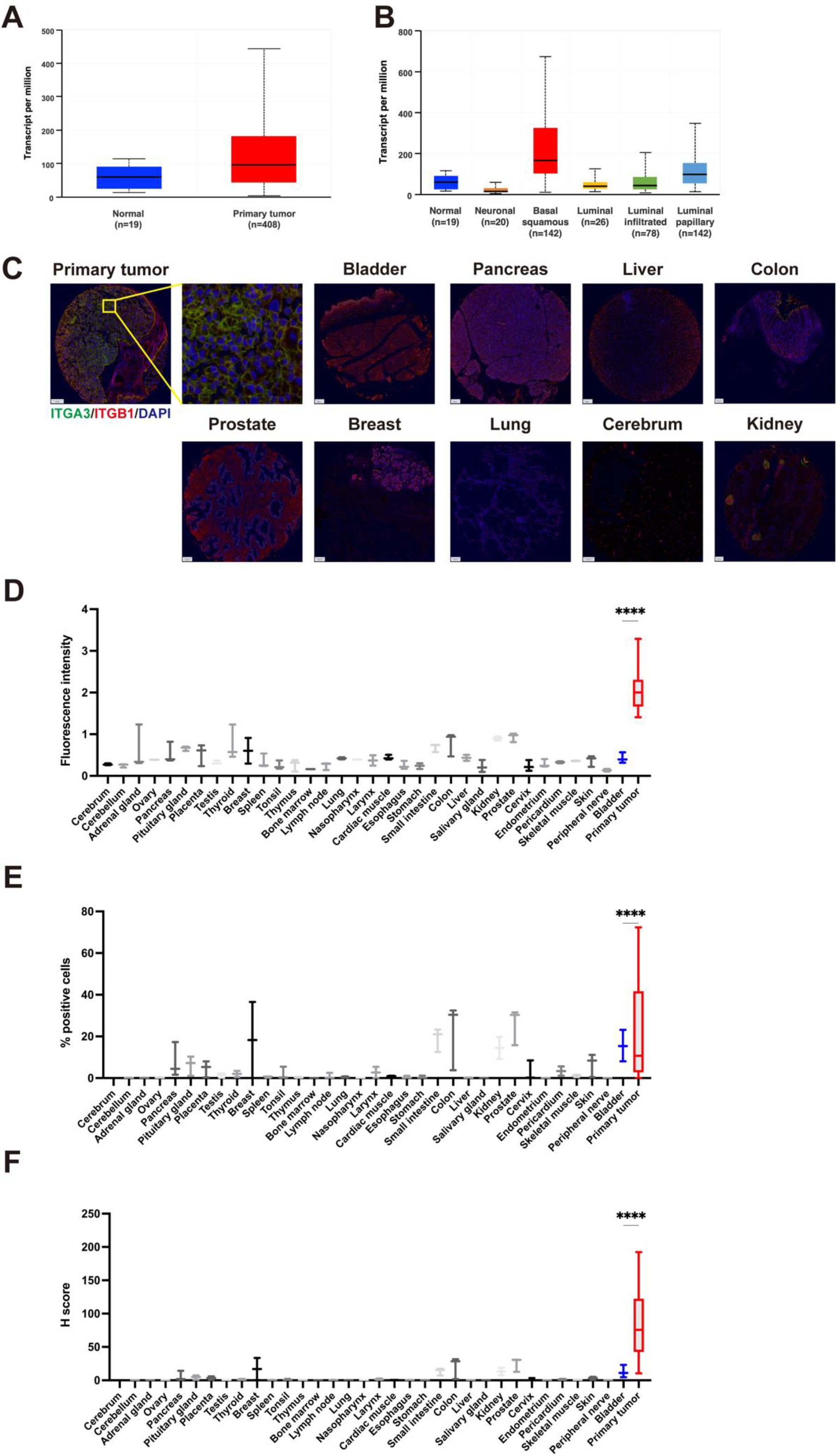
Expression profile of ITGA3 in bladder cancer. **(A)** The expression of ITGA3 was analyzed using TCGA database within UALCAN platform between normal and bladder cancer patients. **(B)** Bar plots showing ITGA3 mRNA expression levels by consensus molecular subtypes in five public clinical cohorts bladder cancer. **(C)** Representative TMA images of individual cores with tumor as well as fluorescent ITGA3 (green), ITGB1 (red), and nuclear DAPI (blue) staining (scale bars, 100 μm; original magnification, 10x). Whole slide images are included in Supplementary Figure S5 and S6. Intensity of ITGA3 staining **(D)**, percentage of cells with ITGA3 expression **(E)**, and H-scores of normal and bladder cancers tissue samples **(F)**.

Building upon these findings, additional insights into the expression patterns of ITGA3 and ITGB1 were gained from tumor specimens of 20 lethal metastatic bladder cancer patients from the University of Washington TAN program (Table S1) (35), alongside 35 normal tissue samples from various human organs including the bladder, pancreas, liver, prostate, breast, lung, and cerebrum, sourced from US Biomax (Table S2). Using multiplexed immunofluorescence (mIF) staining on a clinically and histologically annotated tissue microarray (TMA), we observed elevated levels of ITGA3 and ITGB1 in 28 primary tumor tissues. To visually illustrate these findings, Figure 6C depicts a representative image of tumor tissues from primary bladder cancer patients. The image shows intense green fluorescence for ITGA3 juxtaposed against a red fluorescence background for ITGB1, suggesting their co-expression within the tumor tissues. This visual evidence aligns with the TMA data, highlighting the variation in red fluorescence intensity across normal tissues. The TMA analysis revealed a significant increase in ITGA3 and ITGB1 fluorescence intensity (Figure 6D) and positive cells in patient cores (Figure 6E), with notably higher H-scores (% cells stained × staining intensity, Figure 6F) in these samples compared to those from normal organs, including normal bladder tissues.

Such variation indicates that while ITGA3 expression is predominantly cancer-specific, ITGB1 is ubiquitously expressed across both cancerous and normal tissues (Figure S7). Together, these findings, corroborated by TCGA data analysis, reinforce that ITGA3 is significantly elevated in bladder cancer tissues, underscoring its potential as both a biomarker and a therapeutic target in bladder cancer pathology.

### In vitro cytotoxic evaluation of anti-ITGA3B1 ADC

To evaluate the therapeutic potential of the anti-ITGA3B1 ADC, we performed cytotoxic assays across a diverse panel of human bladder cancer cell lines: UM-UC-3, SCaBER, SW780, CoCaB1, UM-UC-9, and HT-1376. We quantified ITGA3 expression on these cell lines using quantitative flow cytometry (QFACS), which revealed heterogeneous ITGA3B1 antigen expression within each histotype, ranging from 2.1 × 10^5^ to 4.2 × 10^5^ antigens per cell (Table 2). This panel, including an ITGA3 knock-out isogenic line derived from UM-UC-3, served as our in vitro models to assess the cytotoxic efficacy of the ADC. The ADC exhibited potent cytotoxic effects, correlating with the level of antigen expression on the cells, in the order of UM-UC-3 > SW780 > HT-1376 > UM-UC-9 > SCaBER > CoCaB1, with IC50 values ranging from 0.76 to 8.16 nmol/L. Notably, the ADC was effective against cell lines with both high and moderate to low levels of antigen expression. In contrast, cells lacking ITGA3B1 showed negligible response to the ADC, and the unconjugated antibody displayed no cytotoxic activity (Figure 7).

**Table 2.** Summary of ITGA3 expression and cytotoxicity efficacy of ADC.

| Cancer cell line | UM-UC-3 | SW780 | HT-1376 | UM-UC-9 | SCaBER | CoCaB1 | UM-UC-3/ITGA3KO |
| --- | --- | --- | --- | --- | --- | --- | --- |
| QFACS (ABC) | 423,623 | 379,167 | 356,296 | 325,878 | 328,984 | 318,228 | 290 |
| 2E7 ADC cytotoxicity IC<sub>50</sub> (nM) | 0.76 | 0.78 | 0.97 | 1.27 | 1.11 | 1.23 | >100 |

**Figure 7.**
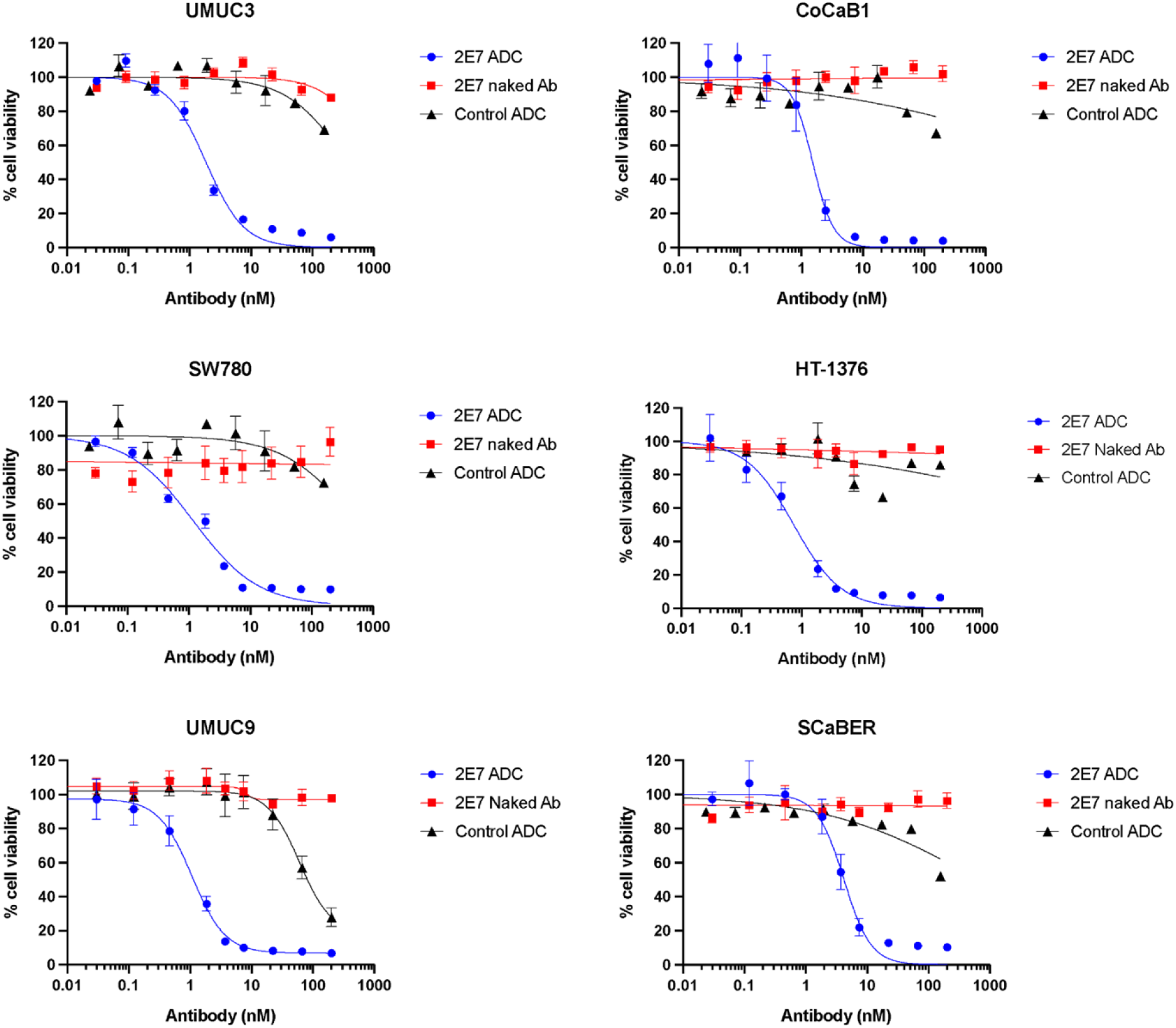
Assessment of cytotoxicity of MMAE-2E7 scFv-Fc ADC on bladder cancer cell lines. The cytotoxicity of MMAE-2E7 scFv-Fc ADC, unconjugated 2E7 scFv-Fc, and a negative control MMAE conjugate with an isotype-matched antibody was evaluated across a range of bladder cancer cell lines. Cell viability was assessed after 72 hours of drug exposure to determine the cytotoxic effects. The cytotoxicity responses measured in terms of IC50 values and percent inhibition.

### Anti-tumor activity of anti-ITGA3B1 ADC in in vivo murine xenograft models

We assessed the therapeutic efficacy of anti-ITGA3B1 ADC in NSG mice using xenografts derived from human bladder cancer cell lines. These mice were employed to develop cell line-derived xenograft (CDX) models. Following tumor establishment, mice received intraperitoneal injections of either a high or low dose of the anti-ITGA3B1 ADC, or PBS as a vehicle control. Tumor volume measurements were the primary endpoint to evaluate therapeutic efficacy.

In xenografts derived from the CoCaB1 bladder cancer cell line (51), a complete regression of tumor growth was observed in the group receiving the high dose of ADC (15 mg/kg administered every five days for five doses), with 60% of these mice remaining tumor-free by day 120 (Figure 8B). Similarly, the same dosing regimen significantly inhibited tumor growth in xenografts derived from the UM-UC-3 cell line, known for its aggressive muscle-invasive urothelial carcinoma characteristics (Figure 8C). In the SCaBER cell xenograft model, we explored doses of 3.75, 7.5, and 15 mg/kg, all of which demonstrated dose-dependent efficacy (Figure 8D).

**Figure 8.**
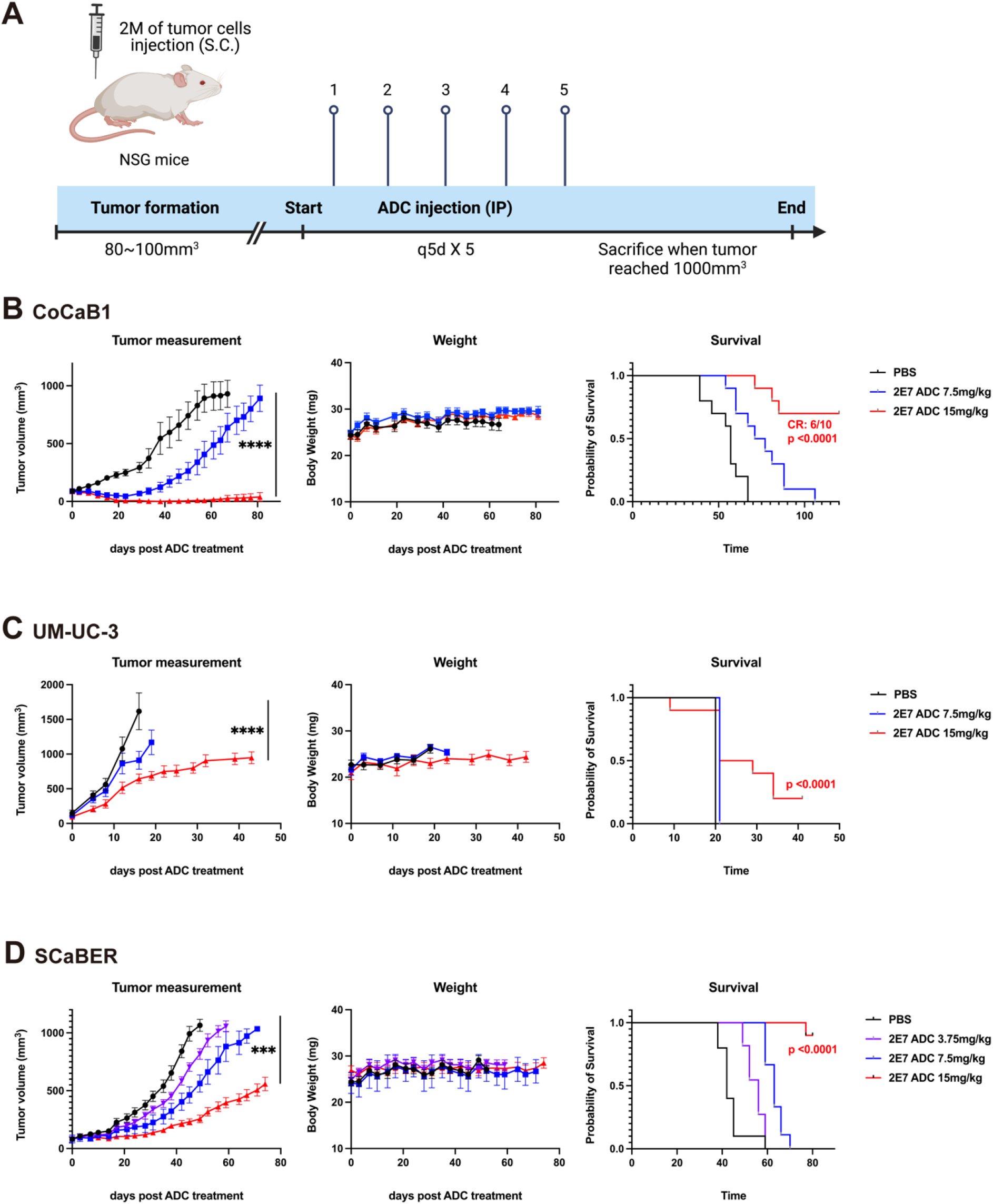
Antitumor activity of MMAE-2E7 scFv-Fc ADC in different sets of bladder cancer xenograft models. **(A)** Schematic and timelines of the in vivo experiments. B-D. Transplanted tumor volumes, body weights and survival rates were evaluated in CoCaB1 **(B)**, UM-UC-3 **(C)**, and SCaBER **(D)** cells. Xenograft-bearing NOG mice with an average tumor size of approximately 100 mm3 received intravenous injections of saline, 7.5 and 15 mg/kg (UMUC3, CoCaB1), or 3.75 mg/kg, 7.5 mg/kg, and 15 mg/kg (SCaBER) every 5 days.

The anti-ITGA3B1 ADC consistently outperformed the vehicle control across all bladder cancer xenograft models tested, achieving significant tumor regression in the CoCaB1 model and robust inhibition of tumor growth in the other models. Importantly, all treatment regimens were well-tolerated, with no observed reductions in body weight or any signs of distress among the treated animals.

These compelling results highlight the effectiveness of targeting ITGA3B1 with an ADC approach in the treatment of bladder cancer. Given the ADC’s capacity to manage different types of bladder cancer effectively, further clinical investigations into ITGA3B1 as a therapeutic target are warranted, potentially opening new avenues for bladder cancer treatment.

## Discussion

Antibody-based therapies, including monoclonal antibodies (mAbs), antibody-drug conjugates (ADCs), and CAR-T cell therapies, have significantly reshaped cancer treatment paradigms (36). Central to their therapeutic success is the identification of Abs with high specificity and functional relevance, which is essential for maximizing efficacy while minimizing off-target toxicity (37). Traditional Ab discovery approaches typically rely on recombinant antigens, which often lack native folding and post-translational modifications. In contrast, phage-display screening on live cells enables the selection of Abs that recognize membrane proteins in their native conformation and cellular context (38), preserving critical structural features and enabling the identification of functionally active Abs, including those with internalization potential (39).

In this study, we advanced this methodology by integrating a target-unbiased live-cell biopanning strategy with an internalization selection step, enabling the discovery of the internalizing Ab clone 2E7. This Ab exhibited efficient receptor-mediated endocytosis, a critical feature for ADCs that depend on intracellular delivery of cytotoxic payloads. To elucidate the cognate antigen of 2E7, we developed an integrative in situ crosslinking-mass spectrometry platform using the amine-reactive crosslinker BS3 to stabilize antibody-antigen interactions on intact cells. This approach circumvents limitations of conventional immunoprecipitation techniques, such as non-specific protein co-isolation and the poor detectability of low-abundance membrane proteins (40).

Through semi-quantitative LC-MS/MS profiling and stringent controls—including counterpart cell lines, isotype IgG, and native cell background—we identified several integrin subunits (ITGA3, ITGA6, ITGA2, ITGAV, ITGB1, and ITGB4) as enriched candidates. Based on known α/β integrin pairings (41), we hypothesized ITGA3B1 and ITGA6B4 as potential heterodimeric targets. Follow-up in vitro and cell-based binding assays confirmed that 2E7 preferentially recognizes the ITGA3B1 complex, displaying markedly enhanced binding compared to individual subunits or alternative heterodimers. These findings establish ITGA3B1 as the functional binding partner of 2E7 and support its candidacy as a novel therapeutic target (42).

Integrins exist in dynamic conformational states regulated by extracellular divalent cations (43–45). In our studies, Mn^2+^ substantially enhanced 2E7 binding to ITGA3B1, whereas EDTA completely abolished the interaction, indicating that 2E7 preferentially binds to the activated integrin conformation (28, 46). Given that elevated Mn^2+^ and Mg^2+^ levels are frequently observed in the tumor microenvironment, this property may enhance the selective activity of 2E7 in vivo (47). This work highlights the value of live-cell biopanning not only for identifying native protein complexes—such as the heterodimeric ITGA3B1—but also for discovering Abs that function optimally under physiologically relevant conditions.

Building on these insights, we explored the therapeutic utility of 2E7 by developing an ADC targeting ITGA3B1. We confirmed ITGA3B1 overexpression in bladder cancer through multiple modalities and noted its strong association with poor prognosis in TCGA data (18), as well as a negative correlation with tumor differentiation scores (50). These findings collectively suggest a critical role for ITGA3B1 in bladder cancer aggressiveness and progression.

To exploit this vulnerability, we generated a 2E7-MMAE ADC, conjugating the 2E7 antibody to the cytotoxic payload monomethyl auristatin E (MMAE), widely used in clinically approved ADCs for its potent bystander-killing effect (30). The 2E7-MMAE ADC showed nanomolar-range cytotoxicity (IC_50_ ∼1 nM) across diverse bladder cancer cell lines and exerted strong, dose-dependent anti-tumor activity in xenograft models. These results establish proof-of-concept for ITGA3B1-targeted ADC therapy in bladder cancer.

Taken together, our study demonstrates a comprehensive platform for phenotypic Ab discovery and target deconvolution using live-cell biopanning combined with in situ proteomics. This integrative approach led to the identification of a novel internalizing Ab targeting the ITGA3B1 complex, with therapeutic application validated through the development of a first-in-class ITGA3B1 ADC. These findings not only support further preclinical advancement of ITGA3B1-directed therapies but also exemplify how phenotypic Ab screening can uncover functionally relevant targets within native cellular environments—an important advancement in the field of targeted cancer therapy.

## Materials and Methods

### 1. Cell lines and culture

HEK-293T, MDA-MB-231, MDA-MB-453 (ATCC) were cultured in DMEM (Gibco) medium supplemented with 10% fetal bovine serum (FBS) and 1% penicillin–streptomycin, unless otherwise stated, at 37 °C, 5% CO2 incubator. CHO-DG44 (ATCC) was maintained in DMEM/F12 (Gibco) supplemented with 10% FBS, 0.1 mM sodium hypoxanthine, 16 μM thymidine, and 100 μg/mL penicillin–streptomycin. UM-UC-3, SCaBER, SW780, UMUC9, and HT1376 (ATCC) were cultured in DMEM (Gibco) medium supplemented with 10% FBS, 1% penicillin-streptomycin and 1% Glutamax (Thermo Fisher). CoCaB-1 was derived from PDX models (51) and cultured in DMEM medium supplemented with 10% FBS, 1% penicillin-streptomycin and 1% Glutamax. Cell lines were passaged in our laboratory for fewer than 6 months after their resuscitation.

### 2. Generation of 2E7 scFv-Fc Antibody

For live-cell panning, 1 × 10^10^ pfu of scFv-phage Abs were incubated with 1 × 10^6^ MDA-MB-453 cells at 4°C for 1.5 hours to allow surface binding. Following incubation, unbound phage particles were removed by centrifugation at 400 × g for 5 minutes and washed with 1% BSA in DMEM. The cell-bound scFv-phage antibodies were then incubated at 37°C for 1 hour in a CO_2_ incubator to induce receptor-mediated internalization. To isolate phage-scFv Abs specific to MDA-MB-453 cells, a non-miscible organic phase separation was performed. Cells were overlaid with an organic lower phase and centrifuged at 10,000 × g for 10 minutes. The lower organic phase was frozen using dry ice, and the supernatant was removed by cutting the tube. The cell pellet was washed with PBS. Surface-bound phage-scFv Abs were eluted using 0.1 M glycine-HCl (pH 2.8) containing 150 mM NaCl. To isolate internalized Abs, cells were lysed using 100 mM triethylamine (TEA). Eluted antibody fragments were analyzed by SDS-PAGE. For flow cytometry, 1 × 10^10^ cfu of scFv-phage Abs were incubated with 1 × 10^5^ MDA-MB-453 cells at 4°C for 1 hour. After washing with FACS buffer (PBS containing 3% BSA and 0.03% NaN_3_, pH 7.2), cells were incubated with mouse anti-M13 phage Ab (1 µg/mL; Thermo Scientific) for 1 hour at 4°C, followed by Alexa Fluor® 647-conjugated anti-mouse IgG (1:800; Jackson ImmunoResearch) for 30 minutes. CHO cells were used as a negative control to assess non-specific binding. After final washing, cells were fixed in 4% paraformaldehyde (pH 7.4) and analyzed using a FACSCalibur flow cytometer with CellQuest Pro software (BD Biosciences).

### 3. LC-MS/MS analysis

4. Proteins were denatured and reduced by boiling at 95 °C for 10 minutes in LDS sample buffer. Samples were resolved by SDS-PAGE using 4–12% gradient Bolt Bis-Tris gels (Invitrogen, MA, USA) at 120 V for 1 hour. Gels were stained with Instant Blue (Sigma-Aldrich, MO, USA), destained with water, and each lane was excised and sectioned into 10 bands using a clean scalpel. Gel pieces (∼1 mm^3^) were destained three times for 20 minutes with 50% acetonitrile (ACN) in 25 mM ammonium bicarbonate (NH4HCO_3_), then dehydrated with 100% ACN and dried in a SpeedVac. Reduction was performed with 25 mM dithiothreitol (DTT) in 100 mM NH4HCO_3_ at 60 °C for 1 hour, followed by alkylation with 55 mM iodoacetamide (IAA) in 100 mM NH4HCO_3_ in the dark for 45 minutes. Gel pieces were washed alternately with 50% ACN in 25 mM NH4HCO_3_ and 100 mM NH4HCO_3_ to remove residual reagents. After drying, proteins were digested in-gel with trypsin overnight at 37 °C. Peptides were extracted sequentially with 10% formic acid (FA), 50% ACN + 0.1% FA, and 80% ACN + 0.1% FA. Combined extracts were dried in a vacuum concentrator and stored at –20 °C prior to LC-MS/MS analysis. For LC-MS/MS, peptides were resuspended in 30 μL of 0.1% FA (Solvent A), and 3 μL were injected onto a trap column (PepMap RSLC C18, 75 μm × 2 cm, 2 μm; Thermo Fisher Scientific). Peptides were separated on an analytical column (PepMap RSLC C18, 75 μm × 50 cm, 2 μm) using a 90-minute linear gradient of 5–35% Solvent B (0.1% FA in ACN) at 300 nL/min: 0–10 min, 2%; 11–12 min, 5%; 13–67 min, 5–35%; 68–83 min, 70%; 84–90 min, 2% for re-equilibration. Mass spectrometry was performed on a Q-Exactive Orbitrap MS (Thermo Fisher Scientific) coupled to an Ultimate 3000 HPLC system. Electrospray ionization was performed at 2.0 kV, with a capillary temperature of 250 °C. Full MS scans were acquired over 400– 1400 m/z at 70,000 resolution. Data-dependent MS/MS was performed using the top 10 precursor ions per cycle with dynamic exclusion (30 s), 17,500 resolution, and normalized collision energy of 27%.

### 5. Database search and statistical analysis

Raw MS/MS data were analyzed using PEAKS Studio version 8.5 (Bioinformatics Solutions Inc., Waterloo, ON, Canada). Processed peak lists were searched against the Homo sapiens reviewed UniProt database (release 2018.09; 20,931 entries). The precursor and fragment ion mass tolerances were set to 10 ppm and 0.5 Da, respectively. Trypsin was specified as the proteolytic enzyme, allowing up to two missed cleavages. Carbamidomethylation of cysteine (+57.0215 Da) was set as a fixed modification, and oxidation of methionine (+15.9949 Da) was set as a variable modification. Protein identifications were filtered with stringent thresholds: peptide -10logP ≥ 20 and protein - 10logP > 20, with a minimum requirement of two unique peptides per protein. For relative quantitation, a label-free quantification approach based on spectral counts was performed using the PEAKS Label-Free tool. Normalized spectral count data from triplicate analyses of two sample groups were subjected to statistical analysis using the Power Law Global Error Model (PLGEM) implemented in R (version 2.15) via the Bioconductor package (http://www.bioconductor.org). PLGEM was used to assess differential protein expression, calculate statistical significance (p-values), and derive signal-to-noise (STN) ratios for each protein.

### 6. Flow cytometry

For standard flow cytometry, cells were dissociated into single-cell suspensions using Versene-EDTA (Thermo Fisher Scientific), washed once with mAb wash buffer (MW; PBS supplemented with 0.1% FBS and 0.1% sodium azide), and resuspended in 100 μL of MW. Cells were stained with 2E7 scFv-Fc antibody at a concentration of 1 µg/mL for 15 minutes at 4°C. After washing, cells were counterstained with 5 µL FITC-conjugated goat anti-human secondary antibody for 30 minutes at 4°C. Following a final wash, cells were resuspended in MW and analyzed on a Sony SH800 flow cytometer (Sony Biotechnology). Data were processed and analyzed using FlowJo software (version 10; RRID:SCR_008520). Quantitative flow cytometry was performed using the Quantum Simply Cellular (QSC) kit (Bangs Laboratories) to determine antibody binding capacity (ABC). Cells at 80–90% confluence were harvested with Versene (Life Technologies), centrifuged at 300 × g for 5 minutes, and resuspended in FACS buffer (PBS + 3% FBS + 0.03% sodium azide). Cells were counted and adjusted to 2 × 10^6^ cells/mL. Aliquots of 2 × 10^5^ cells in 100 μL were dispensed into a 96-well round-bottom plate and incubated with the primary antibody for 1 hour at 4°C. After washing twice, cells were stained with a FITC-conjugated rabbit anti-ITGA3 antibody (LSBio) diluted 1:100 in FACS buffer for 1 hour at 4°C. Cells were then washed and resuspended in DPBS for acquisition. QSC bead standards were prepared and stained in parallel using the same FITC-conjugated secondary antibody per the manufacturer’s instructions. Beads and stained cells were analyzed using a Sony SH800 cytometer. Fluorescence intensity (geometric mean) was obtained for each bead population, and a standard curve was generated using the QuickCal analysis template provided by the manufacturer. The resulting regression allowed calculation of ABC values (reflecting surface receptor density) for the cell samples, enabling quantitative assessment of antibody binding.

### 7. mIF (TMA staining)

University of Washington bladder cancer tumor-associated neutrophil (TAN) tissue microarrays (TMA) and FDA normal organ TMAs (US Biomax Inc.) were used for mIF analysis. Tissue sections were stained using the Leica BOND Rx automated stainer (Leica Biosystems) with Leica Bond reagents. Antigen retrieval and antibody stripping were performed using Epitope Retrieval Solution 2, followed by rinsing with Bond Wash Solution after each step. A high-stringency wash was applied following secondary and tertiary reagent applications using a high-salt TBST buffer (0.05 mol/L Tris, 0.3 mol/L NaCl, 0.1% Tween-20; pH 7.2–7.6). For all secondary detection steps, Opal Polymer HRP Mouse plus Rabbit (PerkinElmer) was used. Quantification of ITGA3 and ITGB1 expression was performed by generating H-scores from mIF data using the CytoNuclear LC v2.0.6 module in the HALO image analysis software (Indica Labs).

### 8. Antibody internalization assay

MDA-MB-453 cells were seeded onto confocal imaging dishes (SPL) and cultured to 70–80% confluence. Cells were washed twice with 1 mL pre-chilled PBS and incubated on ice for 1 hour with either scFv-Fc or scFv-phage antibodies. After extensive washing to remove unbound antibodies, cells were supplemented with 1 mL of serum-free growth medium and incubated for 0 or 2 hours at 37 °C to allow internalization. Subsequently, cells were washed once with cold PBS and fixed with 4% (w/v) paraformaldehyde for 20 minutes at room temperature, followed by two additional PBS washes. For permeabilization, cells were treated with 0.1% Triton X-100 for 5 minutes and washed twice with PBS. The subcellular localization of scFv-Fc antibodies was visualized using Alexa Fluor® 594-conjugated F(ab’)_2_ anti-human IgG (2.5 μg/mL), while scFv-phage antibodies were detected using mouse anti-M13 phage antibody followed by Alexa Fluor® 594-conjugated F(ab’)_2_ anti-mouse IgG (3 μg/mL). Lysosomal compartments were stained using mouse anti-LAMP-1 (BioLegend) or rabbit anti-LAMP-1 (Abcam), depending on the antibody format. Alexa Fluor® 647-conjugated F(ab’)_2_ anti-mouse IgG or anti-rabbit IgG (1:500, Jackson ImmunoResearch) was used to visualize lysosomes. Images were acquired using a Zeiss LSM 880 confocal microscope (Carl Zeiss, Pleasanton, CA, USA) equipped with a 40× water immersion objective and processed using ZEN 3.1 software (Carl Zeiss).

### 9. SPR

The binding affinity between the 2E7 antibody and the ITGA3B1 recombinant protein (Acrobiosystems, IT1-H52Wc) was assessed using the Octet RED instrument (Sartorius), based on bio-layer interferometry (BLI) principles. The 2E7 antibody (50 nM) was immobilized onto Anti-Human IgG Fc Capture (AHC) biosensors for 1200 seconds. After a 600-second wash in kinetic buffer (DPBS, pH 7.4), the sensors were incubated with 50 nM ITGA3B1 recombinant protein prepared with different cofactors (2 mM Ca^2+^, Mg^2+^, Mn^2+^, or 5 mM EDTA) for 3600 seconds (association phase), followed by a 7200-second dissociation phase in kinetic buffer. Affinity constants were determined by fitting the sensorgrams to a 1:1 binding model using FortéBio Data Analysis Software v8.2.

### 10. ADC generation

The 2E7 scFv-Fc ADC was generated by conjugating monomethyl auristatin E (MMAE) to the 2E7 scFv-Fc antibody via a valine-citrulline (vc) linker, following a previously described protocol (52). Briefly, the antibody was partially reduced at 37°C using an excess of Tris(2-carboxyethyl)phosphine. The reduced antibody was then incubated with vcMMAE (MedChemExpress) on ice to allow conjugation. The reaction was quenched by the addition of N-acetylcysteine. Excess reagents were removed, and buffer exchange was performed using a Slide-A-Lyzer™ 10 kDa molecular weight cutoff dialysis cassette (Thermo Fisher Scientific).

### 11. Drug to Antibody Ratio (DAR) Determination

The drug-to-antibody ratio (DAR) of the 2E7-vcMMAE ADC was determined using a polymeric reversed-phase high-performance liquid chromatography (RP-HPLC) system. Analysis was conducted on a PLRP-S column (2.1 mm I.D. × 15 cm, 8 μm, 1000 Å; Agilent Technologies) connected to a Dionex Ultimate 3000-RSLC system (Dionex, Sunnyvale, CA, USA). Gradient elution was performed at 70 °C with mobile phase A (0.05% TFA in water) and mobile phase B (0.05% TFA in acetonitrile, ACN). The gradient began at 10% B for 10 minutes, ramped to 80% B over 30 minutes, held at 95% B for 10 minutes, returned to 10% B in 1 minute, and re-equilibrated for 13 minutes. The flow rate was set at 0.25 mL/min, and detection was performed at 280 nm. DAR fractions were manually collected upon sample elution and concentrated using a 10K molecular weight cutoff centrifugal filter (Millipore, Billerica, MA, USA). The intact mass of each DAR species was further analyzed using matrix-assisted laser desorption/ionization time-of-flight mass spectrometry (MALDI-TOF MS; Microflex LT, Bruker Daltonics, Bremen, Germany). One microliter of each concentrated DAR fraction was spotted onto a stainless-steel MALDI plate, air-dried, overlaid with 1 µL of a matrix solution (20 µg/µL sinapinic acid in 30% ACN with 0.1% TFA), and air-dried again. Mass spectra were acquired in linear positive mode with a detector gain of 20× and a sampling rate of 0.5 GS/s. A total of 1600 laser shots were averaged per sample, with a frequency of 60 Hz, and spectra were calibrated using a BSA/huIgG mixture to span the 20–160 kDa mass range.

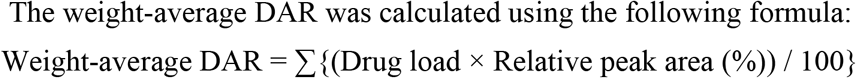

### In vitro cytotoxicity assay

Cells were harvested by trypsinization and seeded into 96-well plates at a density of 4,000–8,000 cells per well. Cells were treated with serial dilutions of 2E7 scFv-Fc-vcMMAE ADC, isotype control ADC, unconjugated 2E7 scFv-Fc, or free MMAE, and incubated at 37 °C in a humidified 5% CO_2_ atmosphere. After 72 hours, cell viability was assessed using AlamarBlue reagent (Life Technologies) according to the manufacturer’s protocol. Fluorescence was measured with a Synergy H1 plate reader (BioTek). Percent viability was calculated relative to untreated control wells. The IC_50_ values were determined by fitting the data to a four-parameter logistic (4PL) nonlinear regression model using the sigmoidal Emax function in GraphPad Prism (GraphPad Software).

### 12. In vivo cytotoxicity assay

All animal procedures were conducted in accordance with protocols approved by the Fred Hutchinson Cancer Center Institutional Animal Care and Use Committee (IACUC) and adhered to Comparative Medicine regulations. Six-week-old male NSG (NOD-SCID-IL2Rγ^null, RRID:BCBC_4142) mice were purchased from The Jackson Laboratory. For tumor establishment, 2 × 10^6^ cells from each human bladder cancer cell line were resuspended in 100 μL of cold Matrigel (Corning) and injected subcutaneously into the flanks of NSG mice. Once tumors reached a volume of 80–100 mm^3^, mice were randomized into treatment groups and received intraperitoneal injections of the indicated agents at specified doses and schedules. Tumor volume, body weight, and body condition scores were assessed biweekly. A complete response was defined as the absence of a palpable or measurable tumor.

## Funding

The National Research Foundation of Korea (NRF) grant RS-2023-00245859 (ECY)

The National Research Foundation of Korea (NRF) grant 2022M3E5F2028363 (KMK)

The National Research Foundation of Korea (NRF) grant 2022R1F1A1074094 (KMK)

The National Research Foundation of Korea(NRF) grant RS-2025-00553130 (KMK)

BK21 FOUR of the Ministry of Education (MOE, Republic of Korea) and NRF (ECY)

The research grant of Kangwon National University in 2022 (KMK)

The research grant of Kyung Hee University in 2024, KHU-202336667 (KPK)

Department of Defense Peer-Reviewed Cancer Research Program awards HT9425-23-1-1055 and HT9425-24-10768 (JKL)

Bladder Cancer Advocacy Network Innovation Award (JKL)

Parker Institute for Cancer Immunotherapy award C-03920 (JKL)

## Author contributions

Conceptualization: ECY, KMK, JKL

Methodology: HRJ, JHL, EHK, YRS

Data Analysis: HRJ, JHL AJS

Investigation: HRJ, JHL, EHK

Resources: ECY, KMK, PSN, JKL, KPK

Visualization: HRJ, MCH

Writing-original draft: HRJ, JHL

Writing-review and editing: ECY, KMK, JKL

Supervision: ECY, KMK, JKL

## Data and materials availability

All data are available in the main text or the supplementary materials. The mass spectrometry proteomics data have been deposited to the ProteomeXchange Consortium (http://proteomecentral.proteomexchange.org) via the PRIDE partner repository with the dataset identifier PXD.

We are thankful for the support of the University of Washington Genitourinary Cancer Research Laboratory and the rapid autopsy team.

